# Open-BEATBox: an open-source autonomous operant chamber for high-throughput data acquisition of complex behaviour

**DOI:** 10.64898/2026.11.07.765109

**Authors:** M. Euvrard, N. Benzina, D. Duarte-Domingues, S.L. Mondragón-González, D. Huzard, G. Penderia, Y. Travert-Jouanneau, O. Lavielle, E. Lousada, C. Schreiweis, A. Lorenz, P. Pavlov, Z. Ajili, L. Lebegue, P. Tissier, E. Burguière

**Author notes:** These authors contributed equally to this work (M. Euvrard, N. Benzina, D. Duarte-Domingues, S.L. Mondragón-González). Current address: Lyon Neuroscience Research Center, Bron, France (E. Burguière).

## Abstract

Behavioural neuroscience increasingly relies on rodent models to investigate the neural mechanisms underlying cognition in health and disease. However, conventional operant testing is typically restricted to short daily sessions that require frequent experimenter intervention and often rely on food or water restriction. These constraints limit behavioural sampling, disrupt natural activity patterns, and introduce variability associated with handling and experimental schedules. Here, we present the Open Platform for Experimental Neuroscience Behavioural and Autonomous Box (Open-BEATBox), an open-source, autonomous home-cage platform that enables continuous, self-paced assessment of complex cognitive behaviour in mice. By allowing animals to live in the testing environment throughout the experiment, Open-BEATBox supports longitudinal behavioural assessment across different cognitive tasks while minimizing experimenter intervention and preserving natural circadian activity. The platform is modular, customizable, and designed to accommodate a broad range of operant paradigms and integration with chronic neural recording and neuromodulation technologies.

Using a visual reversal-learning task, we show that continuous autonomous testing generates approximately ten times more completed trials during reversal learning than conventional session-based protocols while reducing experimenter workload without evidence of major adverse effects on the welfare measures assessed. The flexibility of the platform further enables implementation of additional cognitive paradigms, including uncertainty-based decision-making tasks.

Together, Open-BEATBox provides an accessible open platform for longitudinal cognitive phenotyping, facilitating the collection of large longitudinal behavioural datasets and creating new opportunities to combine continuous behavioural assessment with modern systems neuroscience approaches.

## INTRODUCTION

Behavioural neuroscience aims to understand the neurobiological mechanisms underlying behaviour in both healthy and pathological conditions (Gomez-Marin and Mainen, 2016; Mainen et al., 2016; Niv, 2021). To this end, rodent models have become essential in recent decades, particularly with the development of powerful tools that allow neural activity to be monitored (Siegle et al., 2017; Steinmetz et al., 2018) and manipulated (Park and Carmel, 2016; Whissell et al., 2016) during behavioural tasks. However, major challenges remain in obtaining sufficiently dense and reliable behavioural measures from individual animals while preserving their natural activity patterns and physiological state. This need has contributed to the emergence of home-cage monitoring (HCM) systems, which enables for continuous, automated recording of behaviour directly in the animal’s living environment to reduce the influence of handling, unfamiliar testing environments, and experimenter-imposed schedules (Voikar & Gaburro, 2020; Gaburro & Mandillo, 2026). At the same time, current questions in behavioural neuroscience increasingly require the design of novel complex paradigms capable of modelling, as much as possible, complex sensorimotor or cognitive process observed in humans, often over prolonged training periods (Nithianantharajah et al., 2015). Thus, translating challenging behavioural tasks to mice requires highly flexible experimental apparatuses and often extended training periods.

For this purpose, the development of operant chambers has been widely used to implement complex behavioural tasks adapted from human cognitive paradigms. More recently, the diversity and complexity of learning procedures in rodents have increased with the introduction of visual touchscreen systems (Bussey et al., 2008; Hvoslef-Eide et al., 2016; Mar et al., 2013). However, commercial touchscreen systems can be costly and may offer limited hardware and software flexibility for developing new tasks. As a result, some research groups have developed open-source, low-cost alternatives (Devarakonda et al., 2016; O’Leary et al., 2018; Pineño, 2014; Steurer et al., 2012). In parallel, a range of home-cage monitoring systems (Huzard et al., 2026) has emerged to provide continuous, contactless measurement of spontaneous and operant behaviour in the animal’s living environment. These include sensor-based systems such as the Digital Ventilated Cage (DVC®) (Iannello, 2019), video-based systems such as LiveMouseTracker (de Chaumont, Ey, 2019), automated operant home cages such as AutonoMouse (Erskine et al., 2019) and Souris City (Torquet et al., 2018) and DIY systems based on infrared-red sensors (Virag et al., 2025; Duarte-Domingues et al., 2026). Nevertheless, relatively few platforms combine continuous home-cage monitoring with flexible implementation of longitudinal, cognitively demanding operant tasks. BEATBox was developed to bridge these two approaches by integrating a flexible operant task into a continuously inhabited home-cage environment.

Despite substantial progress in behavioural automation (Schaefer and Claridge-Chang, 2012), many operant studies still rely on discontinuous testing sessions during which animals perform behavioural tasks for periods ranging for 30 minutes to several hours per day (O’Leary et al., 2018; Pineño, 2014). This approach limits behavioural sampling and may introduce variability associated with repeated handling, experimenter presence, motivational procedures, and testing schedules (Balcombe et al., 2004; Vissers et al., 2003). For example, daily handling of animals can be stressful, especially when animals are not adequately habituated (Gouveia and Hurst, 2017; Hurst and West, 2010). It has also been shown that the sex of the experimenter may influence animal behaviour (Neely et al., 2018; Sorge et al., 2014). Another potential limitation of behavioural protocols that depend on the animal motivation is the use of water or food restriction before and during the experimental procedure. It has been shown that food restriction may increase circulating levels of stress hormones (Guarnieri et al., 2012; Levay et al., 2010) and may alter behavioural performance. (Tucci et al., 2006). Finally, the time of the day at which the experiment is run is also a crucial point to consider (Francis et al., 2019). Indeed, rodents are nocturnal animals and many behavioural experiments are conducted during the light phase, which may bias the data and lead to inconsistent results between studies. Together, these limitations; testing nocturnal animals in the light phase, capturing only behavioural snapshots rather than the continuous behavioural activity, and underestimating experimenter and environmental effect, have been framed as the central structural problems that automated home-cage phenotyping is designed to address (Voikar & Gaburro, 2020; Gaburro & Mandillo, 2026).

To address these limitations, we have developed an Open platform: the BEhavioural and AuTonomous Box (Open-BEATBox). Open-BEATBOX is an autonomous home-cage operant platform, in which the animal remains throughout the experiment and self-initiates behavioural trials according to its natural activity and motivational states. The enclosure and task modules are manufactured from laser-cut panels and 3D-printed components, allowing laboratories to reproduce and modify the system using widely available digital fabrication tools. At the core of each Open-BEATBox, a master microcontroller communicates with in-house software running on a host computer and coordinates individual modules, including screens, feeders, sensors and lighting devices. This modular control architecture separates high-level task programming from low-level hardware operation, enabling each box to be reconfigured for different experimental requirements without redesigning the entire system. Multiple boxes can be operated simultaneously by one computer using either identical or distinct task configurations. The main advantage of this autonomous system is that mice can live and move freely in the operant chamber. Thus, the animals can engage in behavioural tasks at their own pace with very few interventions of the experimenter. The automation also guarantees a high throughput data acquisition. This supports both established and novel behavioural protocols, facilitates the use of detailed computational analyses of behaviour, including model-free and model-based approaches, and may reduce variability associated with repeated handling and experimenter-imposed testing schedules.

To validate the Open-BEATBox, we compared continuous 24/7 autonomous testing with a conventional discontinuous protocol in which mice performed a touchscreen-based reversal-learning task for 1 h per day. We assessed task acquisition, trial yield, reversal performance, animal activity patterns and experimenter workload. We show that autonomous continuous testing accelerated progression through successive training stages, generated substantially more trials per animal and reduced the time required for daily experimental supervision, while allowing mice to engage with the task according to their natural activity rhythms

The Open-BEATBox enables more detailed behavioural characterization of individual animals by collecting thousands of trials over prolonged periods in a continuous, self-paced, home-cage environment. By increasing the amount and temporal resolution of data obtained from each subject, the platform may help reduce the numbers of animals required for certain experimental designs, while refining experimental procedures and limiting repeated handling, thereby supporting the principles of the 3RS (Replace the model, Reduce the number of animals, Refine the experimental methods to improve animal welfare) (Richmond, 2000). The continuous acquisition of large behavioural datasets also provides opportunities for future implementation of data-sharing standards aligned with FAIR principles (Wilkinson et al., 2016), which would facilitate data reuse and allowing existing datasets to be reanalysed to address new questions without requiring additional animals (Fuochi et al., 2024; Petit-Demoulière and Huzard, 2026). Finally, the hardware designs, associated software, task protocols and analysis tools are openly available, together with user-friendly interfaces for real-time monitoring and visualization of behavioural performance, with detailed tutorials and step-by-step assembly instructions online (https://open-beatbox.github.io/ and https://github.com/Open-BeatBox). The Open-BEATBox was designed to be built and operated by users without extensive technical expertise. Its open and modular architecture allows the scientific community to adapt, extend and reproduce the system at substantially lower cost than commercial alternatives (White et al., 2019). Together, these features make Open-BEATBox an accessible and complete platform for longitudinal cognitive phenotyping.

## RESULTS

### Open-BEATBox provides a modular, customizable and scalable architecture for autonomous behavioural testing

#### General principles of the modular Open-BEATBox architecture

The Open-BEATBox was designed as a modular behavioural platform that can be readily configured for a wide range of behavioural tasks and with the potential to be adapted to other animal models while remaining compact enough for continuous home-cage experiments and parallel deployment. In the current implementation, the enclosure was carefully optimized for mice resulting in a lightweight compact chamber (363 mm x 363 mm x 300 mm,) that can be easily accommodated within standard animal facilities and laboratories (**Figure 1A**, **left panel**). The Open-BEATBox main structure consists of removable laser-cut panels supporting a variety of interchangeable 3D-printed modules, including feeders, visual displays, and illumination units. Selected modules incorporate flexible buttons directly into the 3D-printed structure. These buttons are created by locally reducing the thickness of the printed material, allowing the plastic to flex when pressed and actuate controls on the underlying electronic board. This design eliminates the need for separate mechanical button components and allows individual controls to be accessed without disassembling the module (**Figure 1A**, **Video S1**). These modules are illustrated here in the context of the reversal-learning task used in this study, which guided the selection of specific components such as touchscreens displays and reward delivery system. (**Figure 1B**). This fabrication strategy allows laboratories to reproduce, repair and modify the platform using widely accessible digital fabrication methods, either through local facilities (e.g. makerspaces or FabLabs) or by outsourcing production to online manufacturing services. Additional task-specific modules can be readily designed and integrated as experimental needs evolve. Detailed designs, assembly instructions and software resources are available through the dedicated website, facilitating adoption and further development by the scientific community: https://open-beatbox.github.io/

**Figure 1.**
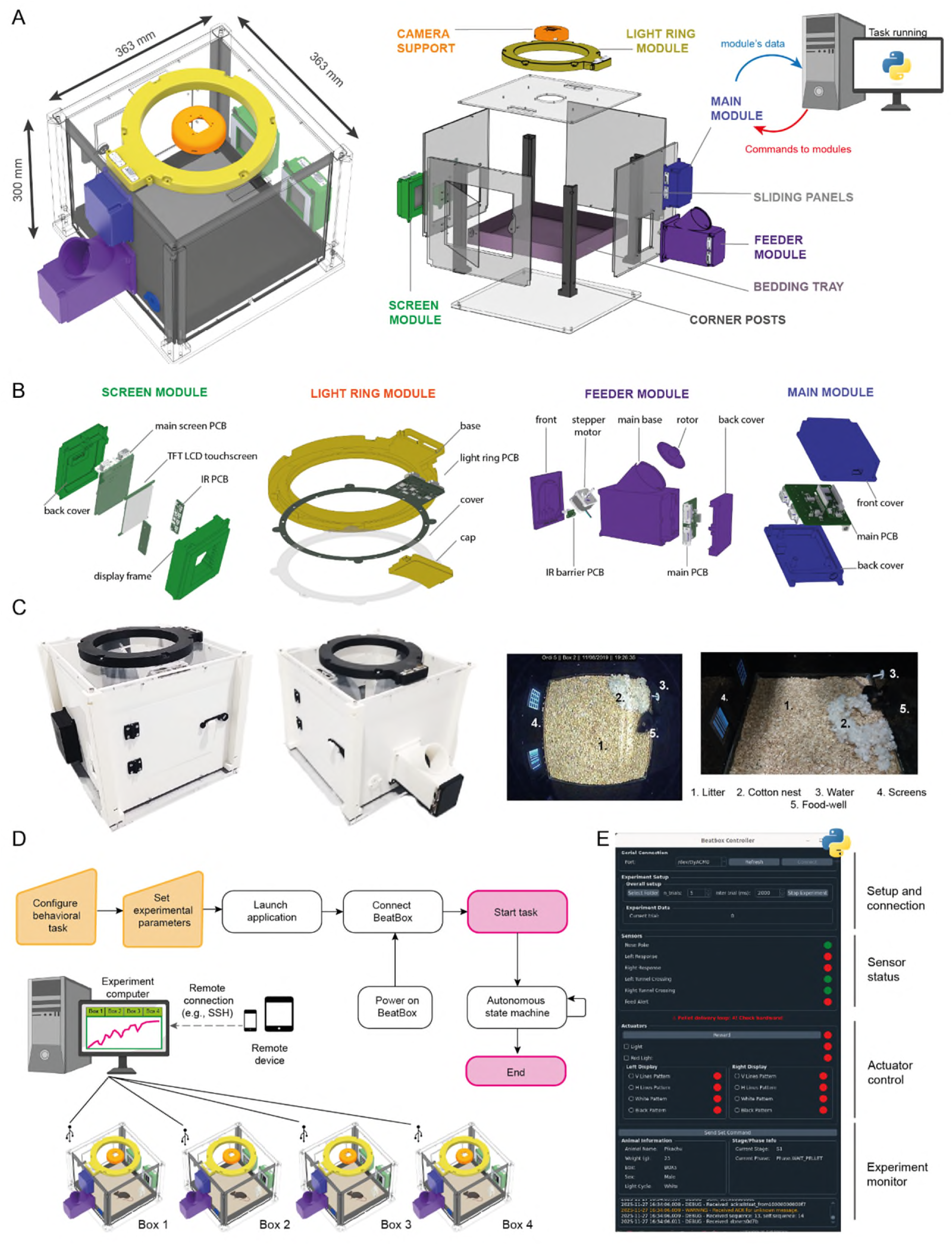
The Open-BEATBox is a highly customizable and user-friendly system. (A) 3D representation of the Open-BEATBox. This operant chamber is highly modular thanks to customizable removable panels and modules. A single computer connected through serial communication to the master RP2040 module is required to run behavioural tasks within the Open-BEATBox. (B) Examples of the 3D-printed / laser-cut modules with all the electronic components used during the study (the Arduino shields are not shown). (C) Pictures of a finalized Open-BEATBox. From the left to the right: front view, back view, top and lateral views of the inside with one mouse performing a trial. (D) The behavioural task and experimental parameters are configured through the PC application. Once the application is connected to Open-BEATBox and the task is started, the task state machine runs in the PC application until completion or manual termination. The dashed lines in the flow chart indicate the parts of the code that can be customized by the experimenter. It is possible to remotely take control of the computer to check the monitoring interfaces and the progress of the mice in the tasks. (E) Monitoring interface to live monitor the mouse activity during the task (top picture) and different parameters of interest (e.g. the performance, bottom picture).

The Open-BEATBox was specifically designed for continuous home-cage operation 24 h/day, 7 d/week allowing the animals to remain in the system throughout multi-week experiments. Importantly, mice can engage in behavioural tasks according to their own natural circadian activity, satiety and motivational states without human intervention. In practice, a mouse can self-initiate a behavioural session by breaking an infrared beam; positioned at the food well or at another task-specific location, thereby triggering a trial. Following a correct response, the mouse can then obtain a food reward and continue to engage with the task according to its motivation and satiety state. Throughout the experiment, animals have *ad libitum* access to water and nesting material and are exposed to the normal light–dark cycle of the animal facility through the transparent roof of the Open-BEATBox (**Figure 1C**). Thus, this autonomous system minimizes experimenter intervention while enabling uninterrupted longitudinal behavioural assessment. Multiple Open-BEATBox systems can be controlled simultaneously from a single computer; with each chamber running either identical or independent behavioural protocols (**Figure 1D**). This scalability allows parallel testing of multiple animals, increases experimental throughput and facilitates the deployment of different behavioural paradigms within the same laboratory. For example, a single experimenter can simultaneously acquire data from multiple animals performing the same behavioural task, thereby accelerating large-scale studies, or conduct pilot experiments using different behavioural protocols across different Open-BEATBoxes in parallel, making more efficient use of the available equipment.

#### Modular and customizable electronic architecture

Precise and reliable timing is critical in behavioural and neurophysiological experiments, where accurate control of stimulus presentation, behavioural events, reward delivery, and their synchronization with neural recordings or neuromodulation is essential for interpreting experimental outcomes. To meet these requirements, particular attention was devoted to the electronic architecture and communication system of the Open-BEATBox. Rather than relying on conventional centralized control architectures, the system implements a dedicated distributed architecture based on CAN bus communication, providing reliable communication between independently controlled modules. Behavioural tasks and data recording are coordinated by a host computer running the experimental software and a main microcontroller that communicates with the independent modules over the CAN bus. Each module locally manages a dedicated component of the apparatus (e.g. feeder, screens, or sensors), while the host computer executes the behavioural logic and records experimental data forwarded by the main microcontroller (**Figure 1A**, **right panel; Figure 1D**, **right panel**). This architecture separates high-level task programming from time-sensitive, low-level hardware control, while allowing modules to be added, replaced, or customized without modifying the overall software framework.

Regarding the electronic architecture, all modules are based on the RP2040 microcontroller. One advantage of the RP2040 is that firmware can be uploaded via USB by dragging and dropping the compiled firmware file onto the device. Another advantage is that the RP2040 supports C/C++ and MicroPython and benefits from an extensive ecosystem of open-source libraries and projects. The communication between modules is handled by the CAN bus, a widely used and robust communication protocol that is often used in automotive applications. Its main advantages include reliable communication, resistance to electrical interference and the ability to connect multiple modules on a shared bus. The controller for this bus is the MCP25625-E/SS, a widely available CAN controller.

All modules share the same core electronic architecture, including the microcontroller and CAN bus controller except for the components required for their specific functions (screen controls, motor control, power distribution, etc.) This common design facilitates users familiar with electronics the development and integration of new modules without requiring substantial modifications to the existing electronic architecture.

#### Modular software architecture and communication protocols

The codebase for both the host computer and the Open-BEATBox modules is implemented in Python. Each module incorporates an RP2040 microcontroller unit (MCU) running MicroPython, a lightweight Python variant optimized for resource-constrained devices. The use of MicroPython facilitates firmware adaptation to additional hardware platforms and the development of new modules.

The system employs two primary communication protocols

● Communication protocol between the host computer and the main module (e.g., USB/UART) for PC-Open-BEATBox interaction.
● An inter-module protocol (e.g., CAN bus) that standardizes packet formats, command structures, and error handling between the main module and secondary modules.

New modules can be integrated by defining the appropriate communication messages and adapting the MicroPython firmware from an existing module template. Together, these design choices enable highly scalable customization, although scalability remains subject to CAN bus bandwidth, message arbitration and hardware limitations.

The Open-BEATBox software is also highly customizable with a main backbone code to initiate the tasks and to ensure the communication between the computer and the main microcontroller. The only requirement for the experimenter is to add the specific functions relative to his / her task that basically consists in the output commands to execute depending on the active inputs (e.g, during the second training stage, if the mouse nosepokes in the food-well, a pellet must be delivered) (**Figure 1D**, **right picture, Figure S2**). Besides, the Open-BEATBox software provides a personalized user-friendly monitoring interface (**Figure 1E**). This interface live-tracks the mice activity all along the task and displays graphic overviews of the evolution over time of any variable of interest (performance, reaction time, lateralization…) (**Figure 1E**). It is possible to directly monitor the experiments from the experimental area or remotely with smartphones thanks to the use of free commercial software or VPN access (**Figure 1D).**

In a benchmark comprising 100,000 pairs of consecutive timestamp calls on an RP2040, the mean measured interval was 18 µs; 36 intervals (0.036%) exceeded 1 ms. Across 9,900 CAN-bus handshake exchanges, no communication failures were observed. Under the default configuration, mean handshake completion times were 22.0 ms for the responding module and 36.2 ms for the initiating module. The round-trip command and acknowledgement exchange between the host computer and a peripheral module took 50.61 ± 3.24 ms (mean ± SD; n = 10,476 exchanges).

### The Open-BEATBox ensures high throughput data acquisition with little experimental effort

To evaluate the Open-BEATBox’s efficiency in collecting large-scale behavioural data, we compared two implementations of the same touchscreen-based reversal-learning task. In the continuous condition, mice remained in the Open-BEATBox around the clock (24 hours a day, 7 days a week), throughout the entire duration of the task and could self-initiate a trial at any time (**Figure 2A**, “Continuous”). This protocol was compared with a discontinuous protocol similar to those commonly described in the literature (Horner et al., 2013; Hvoslef-Eide et al., 2016; Mar et al., 2013; Marquardt et al., 2017; Turner et al., 2017), in which mice performed one hour behavioural session per day in the operant chambers before being returned and isolated to their home cage until the next session (**Figure 2A**, “Discontinuous”). To control for the potential effects of individual housing, two additional control groups were included: group-housed mice (**Figure 2A**, “Group”) and individually housed mice that did not perform any behavioural testing (**Figure 2A**, “Isolation”).

**Figure 2.**
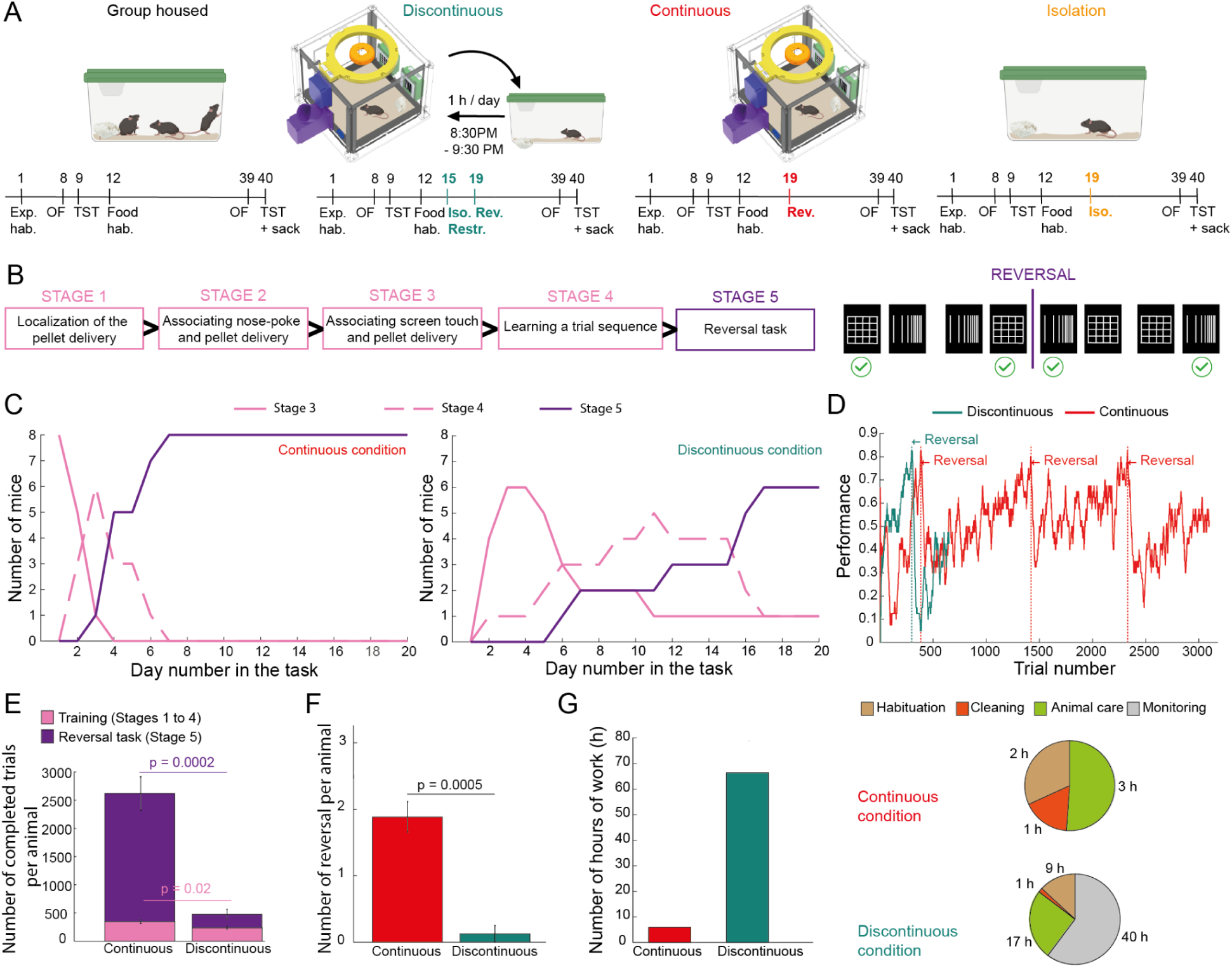
The Open-BEATBox ensures high throughput data acquisition with little experimental effort. **(A)** Presentation of the four conditions used during the protocol and the associated timelines. OF: Open field. TST: Tail suspension Test. Food hab.: Food habituation. Exp hab.: Experimenter habituation. 2 test conditions (discontinuous and continuous) and 2 control conditions (group-housed and isolation) were used. (B) Task protocol. The mice first successively underwent four training stages to get used to the Open-BEATBox and finally reached the stage 5 during which they performed a deterministic stimulus-based reversal learning task (left picture). During this task, the mice had to learn which stimulus from the pair was rewarding and when they reached an 80% performance threshold over forty trials, the contingencies were reversed (right picture). (C) Comparison of the progression in the different stages for the mice in the continuous condition (left) and in the discontinuous condition (right). The mice in the continuous condition reached the reversal learning task faster. (D) Comparison of the performance of two mice (continuous versus discontinuous conditions) during the reversal learning task and localization of the performed reversals (data are smoothed according to a moving average window of 40 trials). (E) Comparison of the number of completed trials during the training stages (stage 1 to 4) and the reversal learning task (stage 5) in the continuous versus discontinuous conditions (n = 8 per condition, data are shown as mean **±** SEM, Wilcoxon rank sum test). (F) Comparison of the number of reversals during the reversal learning task in the continuous versus discontinuous conditions (n = 8 per condition, data are shown as mean **±** SEM, Wilcoxon rank sum test). (G) Comparison of the estimated experimental time in the continuous versus discontinuous conditions. We consider here a total number of 4 mice per condition running the task.

As a representative cognitive paradigm, we selected a touch-screen based deterministic reversal-learning task that has been validated both in the literature (Izquierdo et al., 2017) and in our laboratory using a modified commercial system (Benzina et al., 2021). Following four autoshaping stages allowing animals to progressively learn the task contingencies and get used to the operant chamber (Stages 1-4, **Figure 2B**, **Figure S2**), mice were presented with two visual stimuli (grid on one screen and vertical bars on the other screen) displayed pseudorandomly on two screens (**Video S2, Figure S4**). Responses were detected using infrared break-beam sensors positioned in front of each screen. Animals were required to select the rewarded stimulus, and once they reached a performance criterion of 80% correct responses over the previous 40 trials, the stimulus–reward contingencies were reversed (**Figure 2B**), enabling repeated assessment of cognitive flexibility within the same animal Continuous autonomous testing showed accelerated success in the behavioural acquisition of the training stages compared to the discontinuous condition. After only seven days of training, all mice housed in the Open-BEATBox had reached the reversal-learning phase (Stage 5), whereas only two mice out of the eight mice trained under the discontinuous protocol had reached the same stage on the same day (Figure 2C). Even after 20 days of training, only six of the eight discontinuously trained mice had reached the final stage, demonstrating that unrestricted access to the behavioural task substantially shortened the time required to acquire complex operant behaviours.

All the mice in the continuous protocol completed the different stages of the task within a week, whereas only 6 out of 8 mice reached stage 5 within 20 days in the discontinuous group (**Figure 2C**). The continuous protocol also dramatically increased the amount of behavioural data collected from each animal. This difference is illustrated at the individual level in **Figure 2D**, which shows representative behavioral performance of two mice either in the continuous or the discontinuous protocol. While the mouse in the discontinuous protocol completed one reversal, its paired littermate in the continuous condition completed three, highlighting the advantage of continuous testing in enabling repeated reversal learning over extended periods. At the group-level, mice housed continuously in the Open-BEATBox completed significantly more trials than mice trained under the discontinuous condition (341.75 ± 31.65 trials for the mice in the continuous condition, 231.5 ± 227.80 trials for the mice in the discontinuous condition, p = 0.0207, Wilconxon rank sum test, n = 8 for each condition, **Figure 2E**). This difference became even more pronounced during the reversal-learning stage (Stage 5), where continuous testing generated approximately 10 times more complete trials per animal over the same experimental period (2 272.37 ± 297.12 trials for the mice in the continuous condition, 240.5 ± 88.90 trials for the mice in the discontinuous condition, p = 1.6e-04, Wilconxon rank sum test, n = 8 for each condition, **Figure 2E**). Consequently, mice in the continuous condition (during our 20-day experimental protocol), completed more reversals than those trained under the discontinuous protocol. On average, mice in the continuous condition completed nearly two reversals per animal, while only one mouse in the discontinuous condition reached criterion at all (1.875 ± 0.2266 reversal for the mice in the continuous condition, 0.1250 ± 0.1250 reversal for the mice in the discontinuous condition, p = 4.7e-04, Wilconxon rank sum test, n = 8 for each condition, **Figure 2F**). This increase in data collection in the continuous condition was achieved despite a 10 times reduction in experimenter time, made possible by the fully automated protocol and the minimal need for daily monitoring (**Figure 2G**). Time estimates were based on the time spent by the experimenter each day throughout the procedure. Together, these results show that the continuous Open-BEATBox protocol generates more behavioral data while substantially reducing the practical burden on the experimenter.

### Body-weight stabilization and sustained behavioural engagement during continuous Open-BEATBox testing

To determine whether continuous housing and autonomous behavioural testing in the Open-BEATBox affected animal well-being, we monitored daily body weight, behavioural engagement, circadian activity and standard measures of anxiety- and depression-like behaviours throughout the experimental protocol. An abrupt drop of weight is observed in the discontinuous condition where mice need to be food restricted before the protocol. Mice that underwent the continuous condition also showed a reduction in body weight but more progressively as they reached the most demanding stages of the task (Stages 4 and 5), then their body weight stabilized and remained at or above 85% of their initial free-feeding weight throughout the remainder of the experiment (**Figure 3A**, **3B**). This stabilization indicates that mice maintained a stable energetic state as reflected by their involvement in the task according to their metabolic need. We further examined the relationship between body weight and behavioural engagement by correlating daily body weight with the number of completed trials. For some mice in the continuous condition, there was a significant relationship between the relative mass and the number of performed trials. In three of the eight mice housed continuously in the Open-BEATBox, lower relative body weight was associated with a higher number of completed trials (**Figure 3B right, Table S1 left**). Except for one mouse, all regression slopes are negative, indicating that lower body weight tended to be associated with a greater number of completed trials whereas the opposite tendency was observed in the discontinuous condition (**Figure 3B left, Table S1 right**). Although this relationship was not consistently observed across all animals, these data suggest that behavioural engagement may vary with body weight under continuous testing conditions.

**Figure 3.**
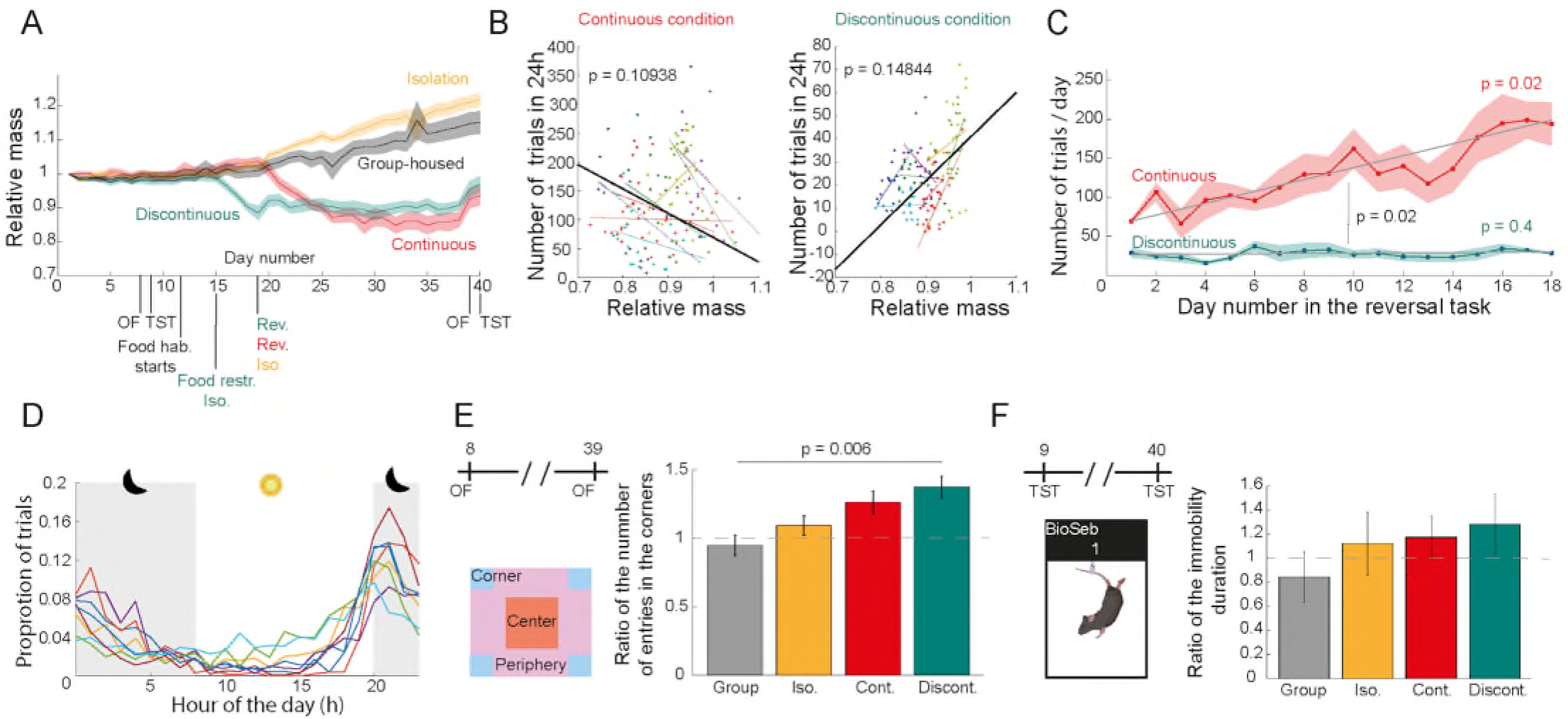
The Open-BEATBox respects the natural phenotype of the animals. (A) Evolution of the body weight for the mice in the different conditions as a ratio with their initial body weights on day 1 (n = 8 per condition, data are shown as mean **±** SEM). (B) Evaluation of the mice activity measured by the number of performed trials every 24h depending on their relative weight. Each colour represents one mouse in the continuous condition (left panel, n = 8) and its paired littermate in the discontinuous condition (right panel, n = 8). The dots represent one measure for each day and the lines represent the polynomial fit (degree 1) for each mouse. The black lines represent the mean of the individual fits (Wilcoxon rank-sum test on the slopes). (C) Evolution of the number of completed trials per day over the training and the reversal task for the mice in the continuous and discontinuous conditions (n = 8 per condition, data are shown as mean **±** SEM, Wilcoxon rank sum test on the slopes after polynomial fitting of degree 1). (D) Distribution of all the trials performed during the reversal learning task along a nycthemeral rhythm for each mouse in the continuous condition. (E) Evaluation of the mice anxiety with the open field test. A ratio superior to 1 indicates that the mice spent more time in the corners after the protocol (post data, acquired on day 39) than before (pre data, acquired on day 8); (n = 8 per condition, data are shown as mean **±** SEM, Kruskal-Wallis test). (F) Evaluation of depressive-like states with the tail suspension test. A ratio superior to 1 indicates that the mice were more immobile and struggled less after the protocol (post data, acquired on day 40) than before (pre data, acquired on day 9). (n = 8 per condition, data are shown as mean **±** SEM, Kruskal-Wallis test).

To further evaluate whether prolonged continuous testing affected behavioural engagement, we quantified the evolution of daily task activity over time. In the continuous condition, the number of completed trials per day, as a proxy of the mouse activity, progressively increased throughout the experiment (mean slope of activity, 6.9227 ± 1.5454, p = 0.0156, Wilcoxon sign rank test for the slopes of all the n = 8 mice, **Figure 3C**, **Table S2 left for individual data**) whereas mice trained under the discontinuous condition maintained a stable level of activity (mean slope of activity, 0.0461 ± 0.3648, p = 0.7422, Wilcoxon sign rank test for the slopes of all the n = 8 mice, **Figure 3C**, **Table S2 right for individual data**). Remarkably, mice in the continuous condition showed sustained and even increased engagement with the task after two weeks of continuous testing, suggesting that the Open-BEATBox maintains motivational drive over extended periods. Mice in both conditions maintained normal levels of task engagement throughout the protocol, with no signs of apathetic behavior despite the weight restriction, single housing, and task training (**Figure 3C**).

One of the principal advantages of continuous Open-BEATBox home-cage testing is that animals can perform behavioural tasks according to their natural circadian rhythm rather than at experimenter-imposed time points. Consistent with this, mice performed most behavioural trials during the dark phase while remaining free to self-initiate trials throughout the 24-hour cycle (Figure 3D, Figure S5A) (**Figure 3D**, **Figure S5A**). Thus, the Open-BEATBox enables continuous assessment of complex behaviour under conditions that better reflect the animals’ natural patterns of activity.

To further investigate the animal wellness, we evaluated anxiety and depressed-like behaviours with the open-field test and tail suspension test before and after completion of the behavioural protocol. We noticed that the mice in the discontinuous condition tended to enter more the open field corners after the 20-day reversal learning task than the mice in the ‘group housing’ control (ratio of the number of entries in the corner after the protocol versus before the protocol, 0.9464 ± 0.0747 for the mice in the group condition, 1.3739 ± 0.0785 for the mice in the discontinuous condition, p = 0.006, Kruskal Wallis test followed by a multicomparison test, n = 8 mice per each condition), suggesting that a discontinuous protocol might induce a significant anxiety (**Figure 3E**). However, we did not find this tendency when looking at the other parameters of the open field (number of entries, time and distance travelled in the center). All the groups did not differ in the tail suspension test (p = 0.4722, Kruskal Wallis test, n = 8 for each condition, **Figure 3F**, **Figure S4**).

Together, these findings indicate that continuous autonomous testing in the Open-BEATBox maintains sustained behavioural engagement, allows mice to perform cognitive tasks according to their natural circadian rhythm, and does not produce strong adverse effects on animal welfare compared with conventional discontinuous behavioural testing.

### The Open-BEATBox allows the implementation of a large variety of innovative tasks

To demonstrate the versatility of the Open-BEATBox beyond the reversal-learning task used for validation, we present several behavioural paradigms that have subsequently been implemented using the same hardware and software architecture in independent projects. These examples illustrate how the modular design of the platform can be readily adapted to different experimental questions without modifying the overall framework (**Figure 4A**).

**Figure 4.**
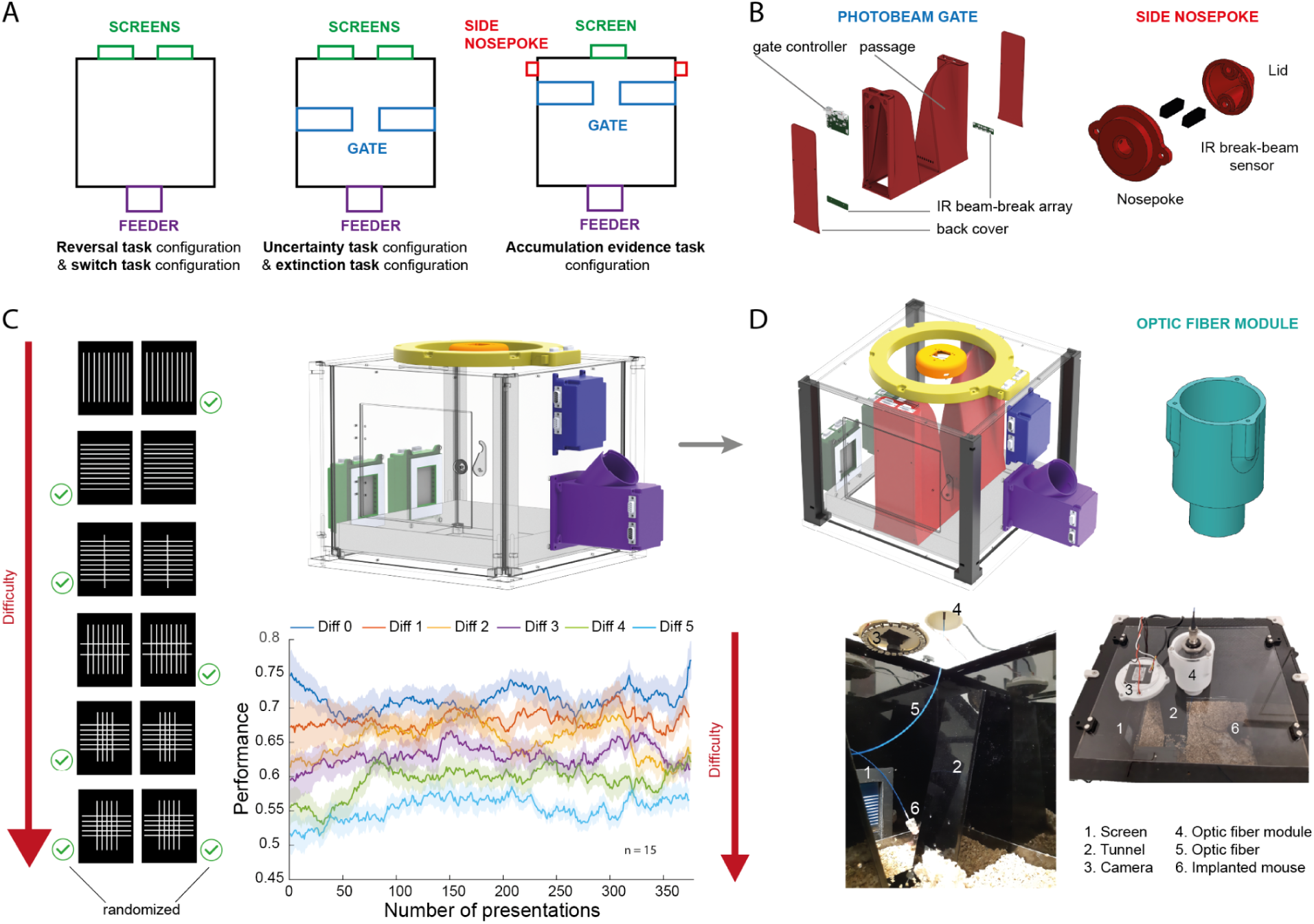
The Open-BEATBox allows the implementation of a large variety of innovative tasks. **(A)** Examples of the three main configurations of the Open-BEATBox used in our team for 5 different behavioural tasks (schemes of the top views). **(B)** 3D-printed / laser-cut tunnels and side nosepoke with the infrared break-beam sensors. **(C)** Example of the uncertainty task developed by Schreiweis and collaborators (Barc et al., 2023). Stimuli with a various ratio of horizontal and vertical bars are displayed on the two screens once the mouse has crossed the tunnel. The mouse has to answer on the left screen if there are more horizontal bars (and vice-versa). The more ambiguous the stimulus (ratio tends to 1), the lower the performance (n = 15 mice, data are shown as mean **±** SEM). **(D)** The Open-BEATBox has been designed to allow the recording and modulation of neuronal activity during behavioural tasks. We show here the Open-BEATBox adapted to calcium imaging with a specific optic fiber module.

In addition to established paradigms such as reversal learning and extinction, three behavioural tasks were specifically developed by extending the existing hardware with task-specific modules (Figure 4A). For example, to improve the temporal precision of reaction time measurements, we designed a tunnel module equipped with two infrared break-beams sensors that continuously track the mouse position within the operant chamber (**Figure 4A**, **middle and right panel, Figure 4B**, **left**). Crossing the tunnel automatically initiates a trial by activating the visual displays, providing a well-defined temporal reference for reaction time measurements compared with tasks initiated by a nose poke in the food well. In another configuration, a side-nosepoke module was designed to collect behavioural responses while presenting stimuli on a unique screen (**Figure 4B**, **right**). These examples illustrate how new behavioural paradigms can be implemented by incorporating dedicated modules while preserving the same hardware architecture and software backbone

Our innovative and flexible set-up was essential to develop these new complex behavioural paradigms that needs extensive training to collect enough data (**Figure 4C**). As a representative example, we implemented an uncertainty task where mice were exposed to a set of 6 stimuli with different ratios of horizontal and vertical bars (Barc et al., 2023). Animals were required to report whether the stimulus contained predominantly horizontal or vertical bars by selecting the corresponding response on the screen. Over a short three-month period for the entire experiment, fifteen mice completed approximately 400 trials for each difficulty level, corresponding to approximately 2 400 trials per mouse. As expected, behavioural performance progressively approached chance level as stimulus ambiguity increased, demonstrating the necessity to collect large numbers of trials per difficulty levels in such perceptual decision-making paradigms (**Figure 4C**).

The modular architecture also facilitates the integration of additional experimental modalities. In an extended version of the uncertainty task, we developed a new configuration combining a single display, dual side nose-pokes and an optical fibre module for simultaneous calcium imaging during behaviour (**Figure 4D**). The same modular framework is readily compatible with electrophysiological recordings, fibre photometry and neuromodulation approaches through dedicated synchronization and stimulation modules, enabling behavioural experiments to be combined with contemporary systems neuroscience techniques.

Collectively, these proof-of-concept implementations demonstrate that Open-BEATBox is not restricted to a single behavioural assay but provides a flexible and extensible experimental platform that can be rapidly adapted to diverse cognitive paradigms and integrated with neural recording and neuromodulation technologies while preserving the advantages of continuous autonomous behavioural testing.

## DISCUSSION

### The Open-BEATBox is an open-source, modular and cost-effective platform for autonomous behavioural testing r

With the Open-BEATBox, we present a cost-effective, automated operant chamber designed for flexible and customizable behavioural testing (**Figure 1**, **Figure 4**, **Video S1**). Our open-source system addresses the growing need for accessible research tools that facilitate reproducibility, transparency and collaboration across laboratories (White et al., 2019). The Open-BEATBox also offers several advantages associated with automated behavioural testing (Schaefer and Claridge-Chang, 2012). The modular architecture allows existing behavioural paradigms to be adapted and new tasks to be developed (**Figure 4**). Providing open-source and sharable hardware and software may also improve standardization across laboratories in the scientific community as many teams will be able to use the same hardware and software configurations, potentially reducing variability associated with differences in experimental equipment. Beyond sharing hardware and code, the high-throughput continuous output of the Open-BEATBox makes it well suited to FAIR data-sharing practices (Wilkinson et al., 2016; Petit-Demoulière and Huzard, 2026). The broader home-cage monitoring field is converging on standardised data formats, shared ontologies, APIs and minimal metadata standards to enable cross-platform reuse, multi-centre comparison and meta-analysis (Fuochi et al., 2024); Future adoption of these standards could facilitate data sharing, reuse and comparisons across laboratories. The Open-BEATBox also ensures a high-throughput acquisition of multiple, various, and complex behavioural data (reaction time, intertrial interval, pellet recovery time, duration of self-exposure to visual stimuli…) over weeks for multiple animals self-training in parallel, saving time for the experimenter to perform cutting-edge quantitative analysis and modelling during this period (**Figure 2**, **Figure 4**). Besides, the limitation of the involvement of the experimenter has two other main advantages. Reduced handling may benefit animal welfare and limit experimenter-related variability, but increased reproducibility across laboratories has not been tested Minimizing repeated handling may reduce experimenter-related variability and handling-associated stress (Balcombe et al., 2004) (**Figure 3**, **Figure S2**) as experimental intervention is largely limited to routine animal care and system maintenance. The rest of the task can be administered and supervised remotely (**Figure 1D**, **Figure 1E**).

Finally, our Open-BEATBox protocol of continuous, 24 h/day testing allows behavioural activity to be assessed across the full light–dark cycle such as the circadian rhythm (**Figure 3C**, **Figure3D**), the recording of physiological parameters such as neuronal activity (**Figure 4D**) and the possibility to perform experiments at specific times within the light–dark cycle (for example, the administration of pharmacological compounds). This represents a major advantage of continuous home-cage monitoring approaches, where disease- and treatment-related phenotypes can occur in the active (dark) phase that conventional daytime testing misses (Pernold et al., 2023; Tomanelli et al., 2024).

While we validated the Open-BEATBox in mice, its modular design may also facilitate adaptation to other species, although this remains to be validated experimentally,as it has previously been done by Steurer and collaborators using comparable behavioural tasks across species (Steurer et al., 2012). The Open-BEATBox costs around US $900. The required components are widely available, and even if the Open-BEATBox assembly requires laser-cut and 3D-printed components, access to university fabrication facilities, FabLabs or external manufacturing services can facilitate its construction.

Routine maintenance and operation of the Open-BEATBox require relatively limited experimenter intervention. Nevertheless, regular checks and occasional cleaning remain necessary to ensure reliable operation, particularly of the food pellet dispenser, as some pellet formulations tend to generate dust that may interfere with reward delivery.

To obtain a more comprehensive characterization of animal activity, existing open-source tracking systems could be integrated (Matikainen-Ankney et al., 2019). Additionally, adding a system to monitor water intake (Godynyuk et al., 2019) could also provide useful information, notably for pharmacological or pharmacogenetic studies. Sound-proofing the overall Open-BEATBox could also be an interesting improvement to prevent the mice from stressful noises from the animal facility. Such additions would move the Open-BEATBox toward a multimodal home-cage monitoring platform, integrating task performance with continuous locomotion, feeding and drinking, circadian activity measures and environmental parameters.

The Open-BEATBox platform has undergone several technical improvements since its initial development. The behavioural experiments presented in this study were performed using an earlier implementation based on Arduino microcontrollers and MATLAB software. The current open-source version retains the same principles of autonomous behavioural testing while introducing an updated architecture based on RP2040 microcontrollers, MicroPython firmware and CAN-bus communication, together with a Python application running on the host computer. Future developments may also include wireless communication and data acquisition to further expand the experimental possibilities of the platform (Amaral-Júnior et al., 2019; Balzani et al., 2018; Francis et al., 2019). The openly available software includes a configurable implementation of the reversal-learning task, together with the associated analysis scripts. Task parameters, such as performance criteria and the number of reversals, can be adjusted to meet specific experimental requirements. Additional behavioural paradigms are being developed, and we encourage users to share new tasks and analysis tools to support further development of the platform.

## STAR METHODS

### Key Resources Table

| **REAGENT OR RESOURCE** | **SOURCE** | **IDENTIFIER** |
|---|---|---|
| **Experimental Models: Organism/Strain** |  |  |
| Mouse; C57BL6J | Brain and Spine Institute |  |
| **Softwares and Algorithms** |  |  |
| Matlab 2017b | MathWorks | https://fr.mathworks.com/products/matlab.html |
| Arduino 1.8.5 | Arduino | https://www.arduino.cc/ |
| Solidworks | Dassault systèmes | https://www.solidworks.com/fr |
| Open field | CleverSys | http://cleversysinc.com/CleverSysInc/csi_products/open-field/ |
| BIO-TST | Bioseb | https://www.bioseb.com/fr/anxiete-depression/2-tail-suspension-test-wireless.html |

### Lead Contact

Further information and request for resources should be directed to and will be fulfilled by the Lead Contact, Eric Burguière.

### Materials Availability

This study did not generate new unique reagents.

### Data and Code Availability

Data and code supporting main results, and the task software are available at Github: https://github.com/Open-BeatBox

### Open-BEATBox architecture and development

Open-BEATBox was initially developed as an autonomous operant platform based on Arduino microcontrollers and MATLAB software. The platform has since evolved into a fully open-source implementation incorporating RP2040 microcontrollers, MicroPython firmware and a Python-based host application. The current architecture retains the core principles of autonomous, self-paced behavioural testing while introducing a modular electronic design based on CAN-bus communication between the main controller and peripheral modules.

The following sections describe the current Open-BEATBox implementation, including its hardware and software architecture and technical performance. The earlier MATLAB/Arduino implementation used to acquire the behavioural data reported in this study is described separately.

### Open-BEATBox hardware

The Open-BEATBox enclosure consists of a modular structure assembled from laser-cut acrylic panels and interchangeable 3D-printed components (**Figure 1A**, **Video S1**). The main structure comprises a cubic frame with four removable lateral panels and an access door. The dimensions and configuration of the enclosure can be adapted to specific experimental requirements, including the animal model, behavioural paradigm and integration of additional recording or stimulation equipment. Task-specific modules can be mounted onto the lateral panels or incorporated within the chamber, allowing components such as visual displays, feeders and infrared sensors to be added or repositioned without redesigning the entire apparatus (**Figures 1A and 4A**).

The enclosure and modules were designed using SolidWorks (Dassault Systèmes). The panels can be manufactured using laser-cutting equipment, while the interchangeable modules are produced by 3D printing. The modular design also allows additional openings or mounting points to be incorporated into the panels to accommodate task-specific components or neural recording devices.

The electronic architecture is organized around a main controller and independently controlled peripheral modules. Each module incorporates an RP2040 microcontroller running MicroPython firmware and communicates through a CAN bus using an MCP25625-E/SS controller. Communication between the main module and the host computer is established through a serial connection. This distributed architecture allows individual modules to manage their associated hardware components while the host computer coordinates the behavioural task.

The peripheral modules share a common electronic architecture, including the RP2040 microcontroller and CAN-bus interface, while incorporating components specific to their function. This design facilitates the development of additional modules by reusing the existing electronic circuitry and communication framework.

To facilitate hardware customization, the BB_tunnel printed circuit board includes a connector compatible with Adafruit and SparkFun I²C modules, as well as accessible pads connected to unused microcontroller input/output pins. The infrared light-curtain modules can provide either individual beam-interruption signals or a combined output indicating beam interruption, depending on the connector configuration. This allows related circuit designs to be used for different applications, including the detection of responses to visual stimuli and the tracking of mouse movements through a tunnel. Infrared beams modulated at 38 kHz are used to reduce interference from ambient light.

Multiple Open-BEATBox chambers can be connected to a single host computer, allowing independent behavioural protocols to be executed in parallel (**Figure 1D**).

### Open-BEATBox software

The current Open-BEATBox software comprises a Python application running on the host computer and MicroPython firmware running on the RP2040 microcontrollers. The host application manages task configuration, behavioural logic, data acquisition and experimental monitoring, while the peripheral modules execute hardware-specific functions and communicate with the main controller through the CAN bus.

Communication between the host computer and the main module is established through a serial interface. The main module forwards commands to the relevant peripheral modules and relays information on hardware events and module states to the host application. Communication between modules follows a common protocol defining message formats and command structures, allowing additional hardware modules to be integrated within the existing software architecture.

Behavioural paradigms are implemented within a common software framework that separates task-specific logic from hardware communication. Experimenters can define the sequence of behavioural events, response contingencies and reward delivery rules according to the requirements of a given task. For example, a nose-poke detected by an infrared sensor can trigger a visual stimulus or the delivery of a food reward, depending on the current task stage. This organization allows different behavioural paradigms to be implemented without modifying the underlying communication framework.

The software also provides a graphical user interface for configuring experiments and monitoring behavioural activity (**Figure 1D,E**). Experimental parameters can be adjusted before starting a task, while the monitoring interface provides real-time information on task progression, animal responses and selected behavioural variables, including performance and reaction times. Experimenters can monitor multiple chambers independently from the same host computer. Remote access to the host computer through external remote-access software or a virtual private network (VPN) also allows experiments to be monitored without requiring continuous presence in the animal facility.

### Timing performance and communication reliability

The timing performance and communication reliability of the current RP2040-based Open-BEATBox architecture were assessed at three complementary levels.

First, the execution overhead associated with timestamp acquisition was evaluated by measuring the elapsed time between consecutive calls to the “time.monotonic_ns()” function on an RP2040 microcontroller over 100,000 iterations. This measurement was used to characterize the overhead associated with the firmware timing function.

Second, inter-module communication was assessed using a four-message CAN-bus handshake protocol repeatedly exchanged between two RP2040-based modules. The elapsed time required to complete the handshake was measured to characterize communication latency and reliability. A total of 9,900 handshake exchanges were analysed.

Finally, end-to-end communication was evaluated by measuring the interval between a command issued by the Python application on the host computer and the receipt of the corresponding acknowledgement from a peripheral module, following transmission through the main RP2040 controller. This measurement included communication between the host computer and the main module, CAN-bus exchanges with the peripheral module and transmission of the acknowledgement back to the host.

Communication tests were conducted under two configurations: a default configuration using an SPI clock frequency of 100 kHz and a CAN-bus bit rate of 250 kbit/s, and a higher-speed configuration using SPI at 10 MHz and CAN at 1 Mbit/s. Repeated measurements were used to characterize communication latency, variability and reliability.

### Hardware and software used for behavioural validation

The behavioural experiments reported in this study were conducted using an earlier implementation of Open-BEATBox, in which an Arduino Mega microcontroller communicated with a host computer running MATLAB R2017b (MathWorks). The main controller coordinated the peripheral components through dedicated Arduino boards and managed the exchange of input states and output commands with the MATLAB application via serial communication.

For the reversal-learning experiments, the chamber was equipped with two visual displays (2.8-inch TFT LCD, ILI9341, Adafruit), each controlled by an Arduino Uno through an 8-bit interface. Behavioural responses were detected using infrared break-beam sensors (3 mm IR Break Beam Sensor, Adafruit) positioned in front of the displays within 3D-printed frames. These sensors registered the animals’ responses independently of the capacitive touchscreen interface.

A NeoPixel ring comprising 24 individually addressable RGB LEDs (Adafruit) was used to deliver a brief white-light stimulus following incorrect responses. The same module provided red illumination for video monitoring during the dark phase. Animal activity could be observed using a miniature camera (700TVL CMOS camera) mounted in the ceiling of the chamber.

Food rewards were delivered through a custom feeder adapted from the Feeding Experimentation Device (FED; Nguyen et al., 2016). The feeder consisted of a pellet reservoir and a rotating dispensing mechanism driven by a stepper motor (14HM11-0404S), controlled by an Arduino Uno and an Adafruit Motor Shield V2. An infrared break-beam sensor positioned above the food well detected nose-poke responses, while an additional sensor below the dispensing mechanism monitored pellet delivery (Figure 1B).

The chamber was constructed from opaque panels to limit visual exposure to the surrounding environment, with a transparent roof allowing animals to remain exposed to the light–dark cycle of the animal facility. The floor contained a removable bedding tray, and nesting material was provided throughout the experiment. Water was available *ad libitum* (**Figure 1C**).

The MATLAB application controlled the sequence of behavioural events and recorded the corresponding responses through continuous serial communication with the master Arduino. Task parameters, including progression criteria, response contingencies and reward delivery, were defined within the experimental software. TThe MATLAB application also provided graphical interfaces for task configuration, behavioural monitoring and recording experimental information (**Figure S1**). These interfaces were used during the behavioural experiments reported here and correspond to the earlier software implementation.

The behavioural procedures and training criteria implemented using this configuration are described in the following sections. The additional adaptive training criteria incorporated into the current Python implementation were introduced after these experiments and were therefore not used to generate the behavioural results reported here.

### Animals

A total of 47 specific pathogen-free (SPF) wild-type male C57BL/6J mice were born, weaned at 21 days, and raised in groups (6 mice per cage, and one cage with 5) in our own animal facility in individually ventilated cages on a 12 h light-dark cycle (lights on at 08:00 AM and off at 8:00 PM) at 21°C and 40% humidity. They had unlimited access to water and food and were provided with nesting material. The experiment started when the animals reached the mean age of 4 months. All the procedures were in accordance with European and French laws and conducted in agreement with institutional guidelines.

We set-up 4 conditions per batch for 4 animals coming from the same cage. In each batch, one mouse underwent the continuous protocol, i.e., when it started the behavioural task, it remained constantly in the cage with permanent access to the task until the experimenter stopped the protocol (Continuous). Another mouse underwent a discontinuous protocol as described in the literature. This mouse was isolated and food-restricted to 85% of its initial weight. It performed the task for 1 hour each day during the dark phase (8:30 PM – 9:30 PM), (Discontinuous). We used a socially isolated mouse (Isolation) and a group-housed mouse (Group-housed) that did not perform any task to control for the isolation parameter (**Figure 2A**).

During the whole experiment, all the mice had access to the same food (LabTab AIN-76A Rodent 20 mg, Test Diet), to nesting materials and the litter was changed every week or as soon as the mice were isolated or started the tasks. They had unlimited access to water. All along the tests and procedures, the same female experimenter handled the animals and she always picked them up with open hands (Hurst and West, 2010). The weight was monitored every day, the nest score regularly assessed when changing the cage and we had access to the food intake for all the isolated mice. The overall wellness of the animals was assessed by monitoring the skin turgor, and looking at the animal activity, posture, face, fur cleanness and any sign of distress would have led to the animal exclusion of the study (Spangenberg and Keeling, 2016). None of them reached a limit point.

The experiments were conducted according to a precise schedule (**Figure 2A**). The animals first got used to the experimenter with a daily interaction for seven days. The habituation protocol occurred between 6:30 PM and 8:30 PM, before the beginning of the dark phase but at a moment during which we know from previous experiments in the lab that the mice are active. The habituation included 10 minutes of observation of the mice in their home cage with the lid open for the mice to get used to the experimenter and her scent; then the experimenter put her gloved hand in the cage for the mice to freely explore it; finally, each mouse was handled for 5 minutes using the cup method (Hurst and West, 2010). They were assessed for anxiety and depression-like behaviours before undergoing the reversal task (or the control housing) after getting used to the food. After the task, the mice were assessed once again for anxiety and depression-like behaviours and sacrificed. All the experiments occurred during the dark phase, around 8:30 PM.

### Open-field test

Anxiety-like behaviour was assessed with the open-field (Crawley, 2007; Seibenhener and Wooten, 2015). The apparatus consisted of a square-shaped arena (5000x500x500mm, length x width x height), made of white and grey Plexiglas ® and illuminated at 40 lux. Subjects were placed facing the wall of the top right corner and could explore the apparatus for 15 min. The open field apparatus was wiped with Aniospray after each mouse to remove odours. The open field was subdivided into a central square (200x200 mm) and 4 corners (100x100 mm) (**Figure 3E**). The following parameters were recorded: number of entries in the central zone, time spent in central zone, number of entries in the corners, time spent in the corners, overall travelled distance in the open-field, distance travelled in the central zone. The data were acquired and analysed through the TopScan software (CleverSys). Mice were habituated to the testing room by bringing their home cage into the testing room for a minimum of one hour prior to the test.

### Tail suspension test

Depression-like behaviours were assessed with the tail suspension test (Powell et al., 2012). During the 6-minute TST, the mice were suspended by their tails from a metal hook (Bioseb, Model: BIO-TST4) using a climbstopper device (Can et al., 2012). The experiment was conducted with a light of 5 lux. The distance between the floor and the tail was about 10 cm. The data (immobility, energy and power) were automatically acquired and analysed through the Bioseb software. Mice were habituated to the testing room by bringing their home cage into the testing room for a minimum of one hour prior to the test.

### Reversal learning Task

The mice in the discontinuous condition were habituated to the testing room by bringing their home cage into the room for a minimum of one hour prior to the test. The mice in the continuous condition were habituated for one hour to the testing room on day 1 and then remained in the Open-BEATBox.

#### Pre-training stage

The pre-training phases were necessary for the mice to get used to the behavioural set-up. They were divided into 4 steps (**Figure 2B**, **Figure S2**). First, the mice had to learn where the pellets were delivered. To achieve that, a pellet was automatically distributed in the food well every minute for 30 times (**Figure S2A**). Then the mice had to nosepoke 30 times in the food well to get a pellet (**Figure S2B**). After that, the mice had to learn how to give an answer. When exploring the cage, if they crossed the infrared break beams before one of the two screens, the screens started blinking and a pellet was automatically delivered. The mice had 15 s to retrieve the reward otherwise the trial was considered as wrong and the countdown was set back to 10 trials to complete anew (**Figure S2C**). Finally, the animals had to understand the global structure of the task. When they nosepoked in the food well, both screens turned with a big white square on and the animal had one minute to give an answer by breaking one infrared beam and they had 15 s to retrieve the reward. The mice had to complete 10 successive complete trials within the imposed delays otherwise the countdown was set back to 10 trials to complete anew (**Figure S2D**). For the mice in the discontinuous task, we moved them from Stage 4 to the reversal task if they had more than 10 rewards during 3 consecutive sessions.

#### Reversal stage

The mice were exposed to a deterministic stimulus-based reversal task (**Figure 2B**, **Figure S3, Video S2)**. After starting the trial by nosepoking in the food-well, the mice were exposed at the same time to two visual stimuli with equivalent brightness (grid and vertical bars) spatially pseudorandomized and they had to pick the rewarded stimulus. If they answered correctly, the screens started blinking and they had 15 s to nosepoke in the food-well to get a pellet. If they gave the wrong answer, a 5 s bright flash was turned on and the following trials were corrective trials, meaning that the locations of the stimuli were repeated until the mouse gave the correct answer as previously described (Mar et al., 2013). When the mice reached a performance of 80 % over the past 40 trials, the contingencies were reversed, and the previous unrewarded stimulus become the new rewarded one (**Figure 2B**). The initial correct stimulus (grid or vertical bars) was balanced across the pairs of mice.

#### Analysis and statistics

All the analysis, statistics and figures of this article were produced with custom-made Matlab codes (R2017b). Due to the small number of animals, we used non-parametric tests. All the analysis codes are provided with extended comments.

For the OF, TST, weight, nest building, number of trials, number of completed trials reversals (trials for which the mice gave an answer), we gathered the values of each variables of interest recorded in the experiments for each animal. The results are plotted as the mean **±** SEM. For the curves fitting, the Matlab polyfit function (degree 1) was used and we calculated the correlation coefficient and the p-value using the corrcoef function (R was then squared and the p-values test the null hypothesis that there is no relationship between the variables). The appropriate tests were used depending on the samples at stake (Wilcoxon sign-rank test for paired samples, Wilconxon rank-sum test for independent variables, Kruskal-Wallis test followed by a multiple comparison test for multiple data sets).

For the performance curves, the correctness of each answer (1 for correct, 0 for wrong) was collected and the overall vector was smoothed with the Matlab smoothdata function, using a moving average of 40 trials (**Figure 2D**) or 50 trials (**Figure 4C**).

The actograms are based on the calculation of an intertrial interval ITI threshold for each mouse above which two trials are considered as belonging to two different burst of activity (red lines). Briefly for each trial n (n > 2), we calculate the following ratio R and studied its distribution. The first value superior to 95^th^ quantile of the distribution is considered as the threshold value.

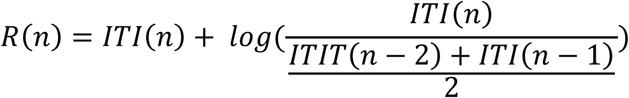

## SUPPLEMENTARY FIGURES AND LEGENDS

**Figure S1.**
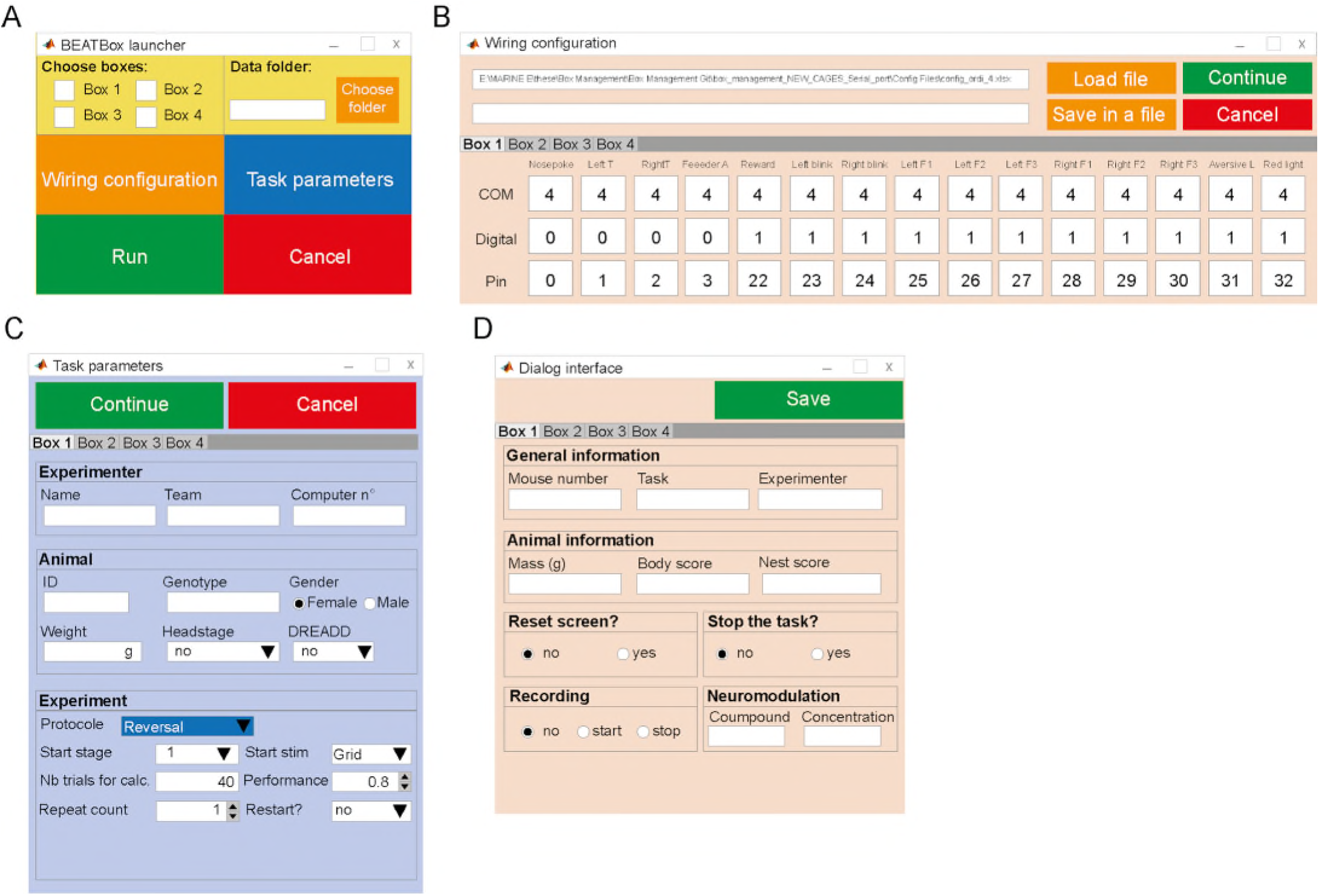
Graphical user interfaces of the original MATLAB-based Open-BEATBox implementation used for behavioural validation. (A) Launcher interface used to configure and initiate behavioural experiments. (B) Interface for assigning hardware input/output addresses and establishing communication between MATLAB and the master Arduino controller.(C) Interface for configuring behavioural task parameters and recording experimental information.(D) Dialog interface used to record animal information and experimental events, including neural recording or neuromodulation procedures.

**Figure S2.**
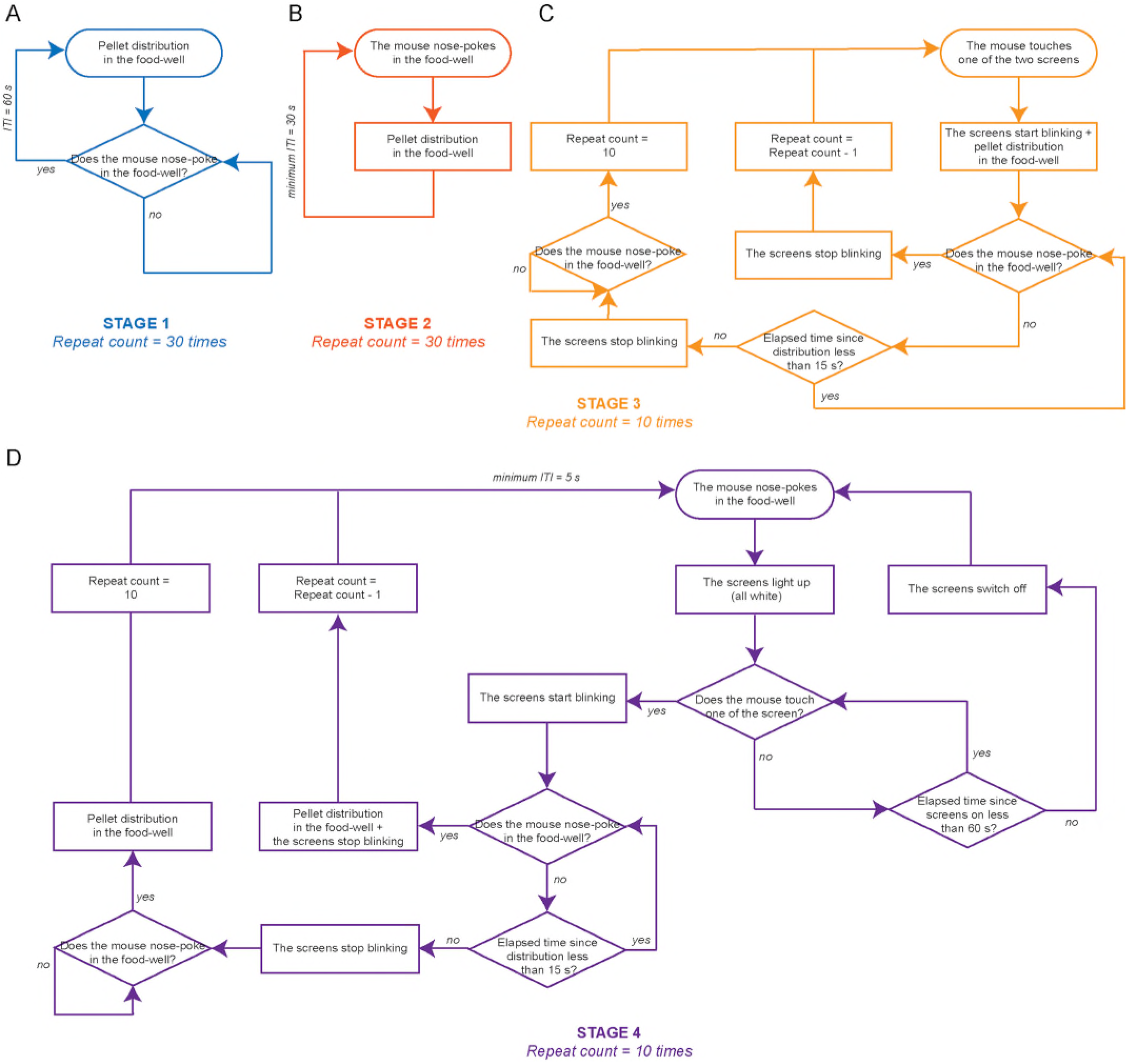
Flow chart of the training stages. Four training stages were required to shape the mouse behaviour towards the final stage. These stages helped the mice to understand how the operant chamber works from the pellet delivery system (Stages 1 and 2) to the answers on the screen (Stage 3) and the way to start and complete a trial (Stage 4). In the most difficult stages (3 and 4) the mice had to complete 10 successive trials (proper actions and timings) in a row to go to the next stages.

**Figure S3.**
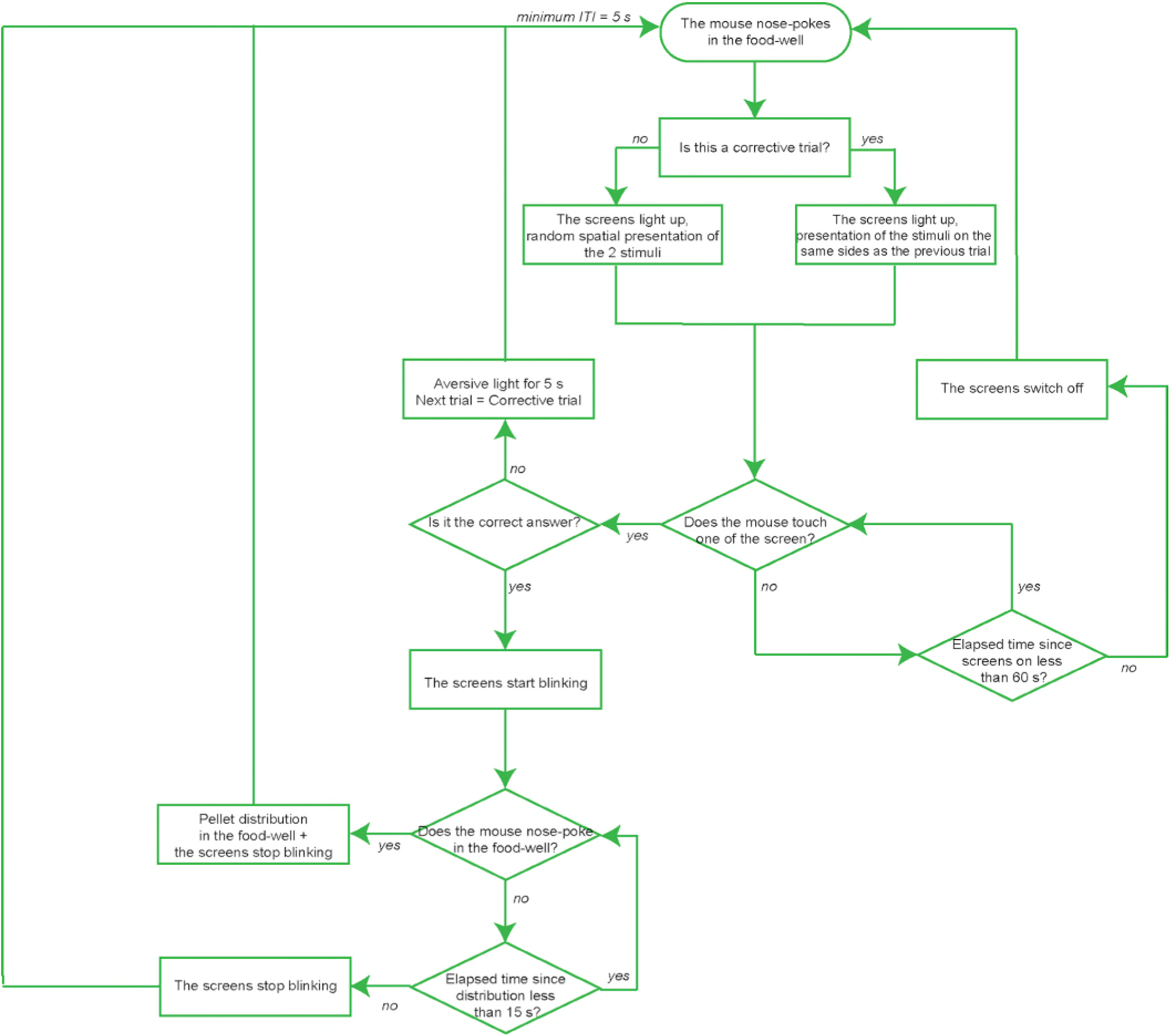
Flow chart of the reversal learning task. During the reversal learning part, the mice had to start a trial by nosepoking in the food-well and pick the side with the rewarded stimulus before 60 seconds. If they gave the correct answer, the screen started to blink, and they had 15 seconds to go to the food-well to get their pellet. If they gave an incorrect answer, a flashlight switched on for 5 seconds. In the latest case, during the next trials the stimuli will be presented on the same sides as the previous trials, otherwise their right/left localisation is random.

**Figure S4.**
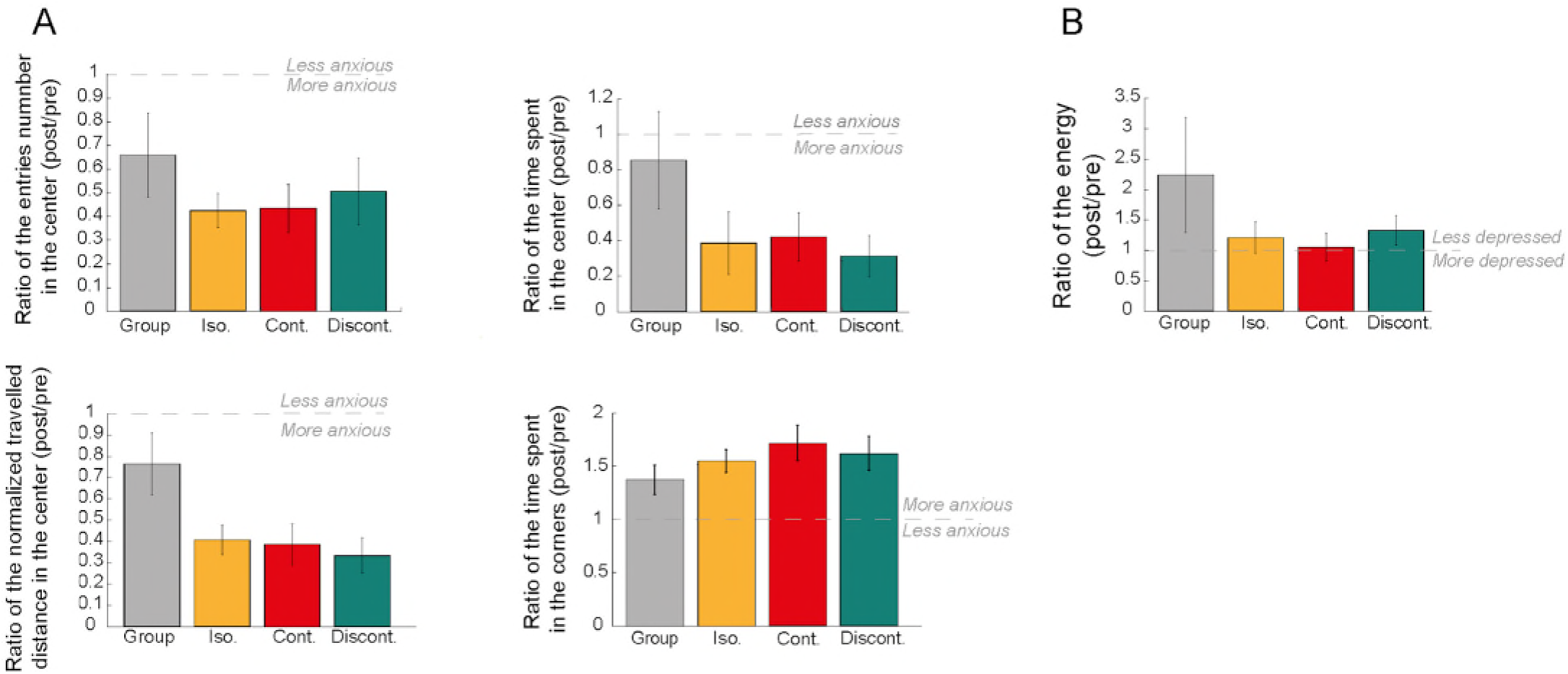
Complementary results of the open field and tail suspension test. (A) Evaluation of the mice anxiety with the open field test. The travelled distance in the center was normalized by the overall distance travelled. For the center data, a ratio inferior to 1 indicates that the mice tend to avoid more the center after the protocol (post data, acquired on day 39) than before (pre data, acquired on day 8) and vice-versa for the corners data. (n = 8 per condition, data are shown as mean **±** SEM, Kruskal-Wallis test, Multiple comparison test if p <= 0.05 in the Kruskal-Wallis test). (B) Evaluation of the mice’s depression-like state with the tail suspension test. A ratio superior to 1 indicates that the mice struggled more after the protocol (post data, acquired on day 40) than before (pre data, acquired on day 9). (n = 8 per condition, data are shown as mean **±** SEM, Kruskal-Wallis test).

**Figure S5.**
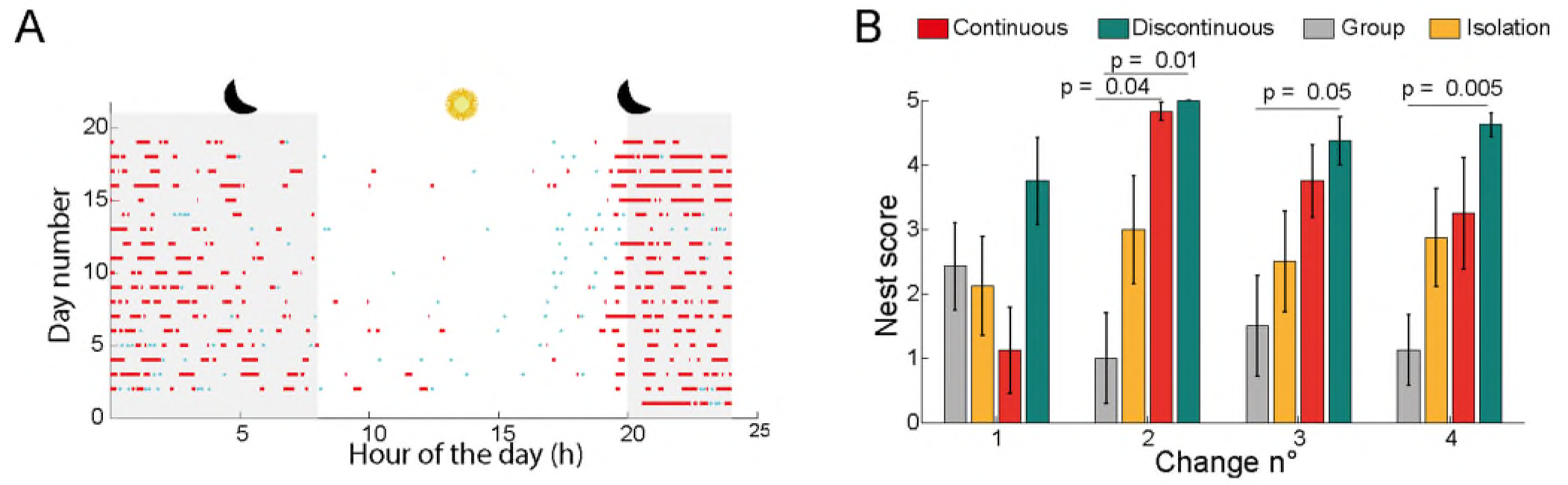
Supplementary assessments of the mice well-being. (A) Example of the activity of one mouse in the continuous condition, along the training and the reversal learning task. Each red line indicates a burst of activity (high number of consecutive trials), the light blue dots indicate isolated trials. (B) After each litter change, a new cotton pad was given to the mice and the nest score was assessed 24h afterwards (Deacon, 2006) (n = 8 per condition, data are shown as mean **±** SEM, Kruskal-Wallis test, Multiple comparison test if p <= 0.05 in the Kruskal-Wallis test).

**Video S1 Flexibility and customizability of the Open-BEATBox**

Each panel and modules are screwable and the Open-BEATBox configuration can be adjusted to the user needs.

**Video S2 Mouse performing the stimulus-based reversal task**

When the mouse nosepokes in the food-well, the screens switch on with the 2 visual stimuli. If the mouse answers correctly, the screens start blinking and the mouse has 15 seconds to get its reward. If the answer is incorrect, there is a strong flashlight. The mouse is being videotaped thanks to the camera from the ceiling and the red light from the Neopixel.

**Table S1.**
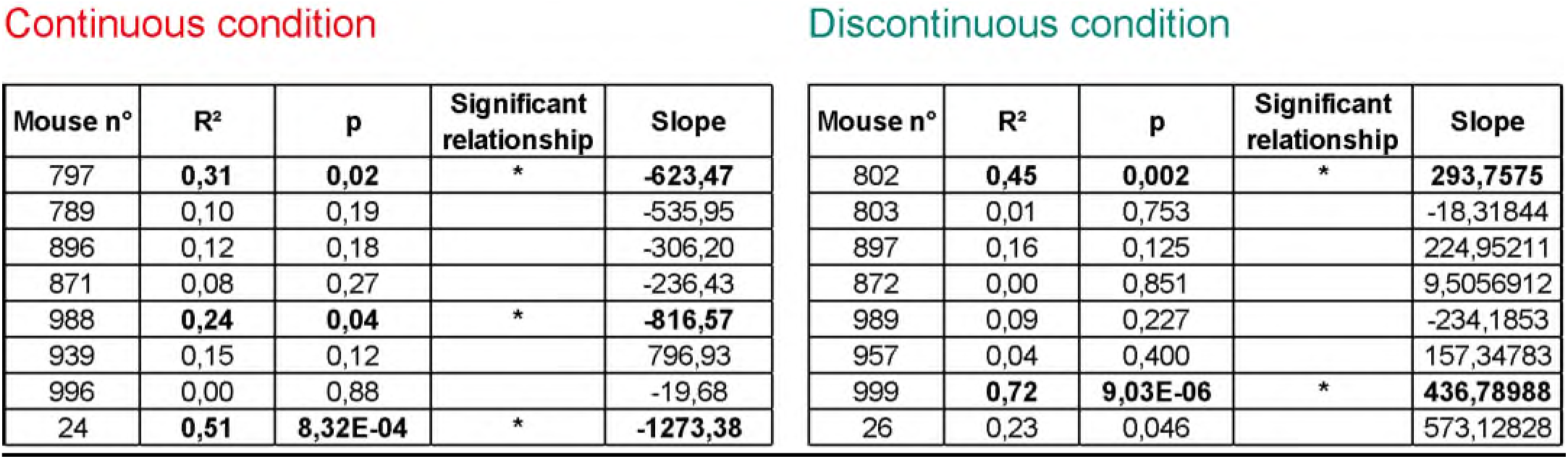
Results (correlation coefficient, p-value and slope) for the individual linear fits of the number of trials against the relative mass of the mice.

| Continuous condition |  |  |  |  | Discontinuous condition |  |  |  |  |
|---|---|---|---|---|---|---|---|---|---|
| Mouse n° | R<sup>2</sup> | p | Significant relationship | Slope | Mouse n° | R<sup>2</sup> | p | Significant relationship | Slope |
| 797 | **0,31** | **0,02** | * | **-623,47** | 802 | **0,45** | **0,002** | * | **293,7575** |
| 789 | 0,10 | 0,19 |  | -535,95 | 803 | 0,01 | 0,753 |  | -18,31844 |
| 896 | 0,12 | 0,18 |  | -306,20 | 897 | 0,16 | 0,125 |  | 224,95211 |
| 871 | 0,08 | 0,27 |  | -236,43 | 872 | 0,00 | 0,851 |  | 9,5056912 |
| 988 | **0,24** | **0,04** | * | **-816,57** | 989 | 0,09 | 0,227 |  | -234,1853 |
| 939 | 0,15 | 0,12 |  | 796,93 | 957 | 0,04 | 0,400 |  | 157,34783 |
| 996 | 0,00 | 0,88 |  | -19,68 | 999 | **0,72** | **9,03E-06** | * | **436,78988** |
| 24 | **0,51** | **8,32E-04** | * | **-1273,38** | 26 | 0,23 | 0,046 |  | 573,12828 |

**Table S2.** Results (correlation coefficient, p-value and slope) for the individual linear fits of the number of trials against the day number in the task.

| Continuous condition |  |  |  |  | Discontinuous condition |  |  |  |  |
|---|---|---|---|---|---|---|---|---|---|
| Mouse n° | R<sup>2</sup> | p | Significant relationship | Slope | Mouse n° | R<sup>2</sup> | p | Significant relationship | Slope |
| 797 | **0,48** | **0,0015** | * | **9,30** | 802 | 0,22 | 0,051 |  | -0,74 |
| 789 | **0,35** | **0,0103** | * | **8,06** | 803 | 0,02 | 0,606 |  | -0,17 |
| 896 | **0,43** | **0,0032** | * | **7,12** | 897 | 0,01 | 0,722 |  | 0,13 |
| 871 | 0,01 | 0,6901 |  | -0,87 | 872 | **0,24** | **0,048** | * | **0,46** |
| 988 | **0,69** | **1,9586E-05** | * | **13,15** | 989 | **0,37** | **0,008** | * | **1,42** |
| 939 | 0,16 | 0,1007 |  | 4,82 | 957 | 0,07 | 0,280 |  | 0,99 |
| 996 | 0,16 | 0,0959 |  | 3,53 | 999 | **0,53** | **0,001** | * | **-1,90** |
| 24 | **0,63** | **0,0001** | * | **10,27** | 26 | 0,01 | 0,776 |  | 0,18 |

## AUTHOR CONTRIBUTIONS

Conceptualization, E.B, N.B, M.E, D.D; Methodology, E.B, N.B, M.E, D.D; Hardware, M.E, P.T, Y.T, M.B. Software, M.E, G.P; Resources, M.E; Investigation, M.E; Formal analysis, M.E; Data curation, M.E, C.S; Writing and editing; M.E, SLMG, D.H, E.B; Online resources: SLMG, DH, Funding acquisition, E.B, C.S.

## DECLARATION OF INTEREST

The authors declare no competing interests.

## ACKNOWLEDGMENTS

This work was funded by: Agence Nationale de la Recherche ANR-16-INSERM-SINREP, ANR-19-ICM-DOPALOOPS, ANR-21-ICM-PREDICTOC (EB) and ANR-24-INSERM-EVOMATIC (CS); Fondation de France (EB); Carnot Foundation (EB). All animal work was conducted at the PHENO-ICMice facility. The Core was supported by 2 “Investissements d’avenir” (ANR-10-IAIHU-06 and ANR-11-INBS-0011-NeurATRIS) and the “Fondation pour la Recherche Médicale”.

## Notes

### Competing Interest Statement

The authors have declared no competing interest.

